# DEVELOPMENT OF PROTOCOLS FOR DEPOSITING A RARE PLANT SPECIES OF THE ROSTOV REGION, *THYMUS CALCAREUS* KLOKOV & DES. – SHOST

**DOI:** 10.1101/2025.03.31.646457

**Authors:** Vasiliy A. Chokheli, Klimentiy A. Tulupov, Anatoly S. Azarov, Mark O. Belyaev, Christina A. Tsymerskaia, Svetlana N. Sushkova

## Abstract

In the course of this study, a protocol for *in vitro* deposition of a rare plant for the Rostov region, *Thymus calcareus*, was developed. Before sterilization, the seeds must be subjected to weekly extreme stratification (−18°C). An effective method for sterilizing primary explants is a mixture of 96% ethanol and 3% hydrogen peroxide in a 1:1 ratio with an exposure time of 7 minutes, followed by three rinses with distilled water. The effective environment for seed initiation is 1/2 MS. The optimal nutrient medium for clonal reproduction of *T.calcareus* has been determined: 1/2 QL. During long-term cultivation of *T.calcareus* on a nutrient medium of 1/2 QL, 100% rhizogenesis of plants was observed. The effect of nutrient media, growth regulators and their concentrations on the multiplication and rhizogenesis of *T.calcareus* explants in vitro has been studied: for multiplication, it is most effective to use a 1/2 QL nutrient medium with the addition of low concentrations of KIN; for rhizogenesis, it is recommended to use QL nutrient media with low concentrations of IAA. The degree of change in the influence of environmental factors, growth regulators and their concentration on *T.calcareus in vitro* and their change over time on the multiplication rates were determined: the nutrient medium (F_fact_ = 4.72), the type of growth regulator (F_fact_ = 7.82) and its concentration (F_fact_ = 4.71) have a significant effect on the multiplication coefficient of explants. The effect of the factors remains almost unchanged for the mineral base and growth regulator, while the concentration of the growth stimulant increases significantly.; A significant effect on the shoot elongation index was found in the growth regulator (F_fact_ = 10.39) and its concentration (F_fact_ = 5.51), the influence of both factors increases

## INTRODUCTION

The problem of reducing the species diversity and gene pool of plants, due to: land desertification, rapid climate change, environmental pollution and the irrational use of natural resources by humans, is becoming increasingly urgent. Plants perform a number of important functions in nature: they are the primary producers and the basis of the ecological pyramid, participate in the circulation of nutrients and various elements, and also help prevent soil erosion. Their role in human life is also important, they are used as a source of food, medicines, materials for construction, textiles, etc.

Conservation of the genetic diversity of rare species *in vitro* is an important addition to current ex situ methods. The formation of banks of calluses, suspensions, and meristems makes it possible to preserve the genetic diversity of a species, as well as to obtain plant material using micropropagation methods for further introduction or reintroduction, as well as the possible use of species with beneficial traits in breeding or mass cultivation. The cryopreservation process, in turn, consists of freezing plant tissues or cultures at -196 °C (Reed et al., 2011; Molkanova et al., 2016).

Thanks to micropropagation methods, rare and endangered plant species can be preserved, as well as fundamental research of these species can be greatly simplified. Obtaining viable planting material *in vitro*, germinating seeds or vegetatively cloning rare plants can be used for the reintroduction and maintenance of the stability of wild populations, as well as the creation of living collections *in vitro* and cryopreservation.

*Thymus* is a genus of the *Lamiaceae* family. All representatives are low-growing creeping semi-shrubs and shrubs with woody stems, occasionally erect, the main feature of which is the presence of essential oil glands. The range of the genus *Thymus* L. covers almost all of Eurasia, with the exception of the tropical part in the southeast, the desert zone, the West Siberian lowlands and most of Kamchatka. Due to the large range of the genus, it is possible to judge its antiquity and, consequently, the strong variability of species. The appearance of representatives of this genus is extremely variable due to the wide area of distribution and different habitus of plants in different periods of ontogenesis. In addition, the genus *Thymus* L. intraspecific polymorphism is characteristic, and often their hybridization can occur at the contact points of different thyme species, which, with sufficient stability and viability of the offspring, can form a new species of hybridogenic origin. All these factors significantly complicate the study of this genus (Gogina, 1990; Mayevsky, 2014).

The secondary metabolites of *Thymus* species are mainly represented by volatile essential oils and non-volatile phenols. In different species, the quantitative and qualitative composition of these compounds varies depending on internal (seasonal and ontogenetic factors) and external (climate, light, soil) characteristics (Gogina, 1990; Sheremetyeva et al., 2017).

Thyme’s high pharmaceutical value is due to its main biologically active components. Due to the high polymorphism of the species, as well as intraspecific genotypic changes, the genus Thymus is of particular interest in the study of its representatives (Buzuk et al., 2012).

Thus, the purpose of this study is to develop a protocol for depositing a representative of the genus *Thymus*.

## MATERIALS AND METHODS

### Sterilization of explants

Seeds were used as biological material for *in vitro* culture initiation of *T.calcareus*. The seeds were provided by the seed bank of the Botanical Garden of the Southern federal university (SFedU). The seed material was collected from plants cultivated in the nursery of rare and endangered plants of the Botanical Garden jf the SFedU in 2023 and stored in dark, dry conditions at room temperature before the experiment.

To select the sterilizing agent of the seed material, three different techniques were tested, selected not only for seed sterilization, but also for their complete purification from secondary metabolites of the parent plant.

Surface sterilization of seeds was carried out using the following sterilizing agents: C_2_H_5_OH (96%), C_2_H_5_OH (70%), H_2_O_2_ (3%). After treatment with sterilizing agents, the seeds were rinsed three times with sterilized distilled water for ten minutes. The sterilization methods used in the study are shown in Table 1.

**Table 1.** Selected methods of sterilization of *T.calcareus* seed explants.

| Method Number | I | II | III |
| --- | --- | --- | --- |
| Sterilizing agents | C<sub>2</sub>H<sub>5</sub>OH (96%) + H<sub>2</sub>O<sub>2</sub> (3%) 1:1 | C<sub>2</sub>H<sub>5</sub>OH 96% | C<sub>2</sub>H<sub>5</sub>OH 70% |
| Exposure time | 7 min | 5 min | 7 min |

The seeds were germinated at a constant temperature of 25 °C with a sixteen–hour photoperiod. After 9 days, the number of infected explants (%) was determined.

### Germination of seeds in vitro

After sterilization, explants under aseptic conditions were inoculated into glass jars (5*7 cm) containing 15-20 ml of Murashige and Skuga nutrient medium (MS) in three different concentrations of micro- and macronutrients with the addition of 3% sucrose and 0.7% agar (MS, 1/2 MS, 1/4 MS). In each jar, 5 seeds were planted 2-3 cm apart; the number of repetitions was fifteen times for each medium.

The seed material, which is not subject to stratification, was collected and cleaned in 2023 and stored in dark, dry conditions at room temperature for several months, another batch of seeds was subjected to extreme stratification for a week at t = -18 °C before sterilization and planting.

The germination of seeds and their anatomical and morphological characteristics of sprouts were monitored for 9 days with an interval of 3 days.

### Multiplication of shoots and rhizogenesis of T.calcareus in vitro

To study the effect of nutrient media on the growth and development of shoots and roots of *T.calcareus in vitro*, apical and lateral shoots obtained by germination of primary explants, 0.8 – 1.5 cm long, were cultured on 6 different variants of hormone-free nutrient media: MS, 1/2 MS, B5, 1/2 B5, QL, 1/2 QL.

Shoots obtained during *in vitro* seed germination were also used to study the effect of growth regulators. Explants were cultured under aseptic conditions on 12 modified variants of MS culture medium with the addition of two types of cytokinins and auxins: BAP (6-Benzylaminopurine) and KIN (kinetin) at concentrations of 0.5, 1 and 2 mg/L; 2,4-D (2,4-Dichlorophenoxyacetic acid) and IAA (Indolyl acetic acid) in concentrations of 0.05, 0.1 and 0.5 mg/L.

Variants of modifications of the MS environment are provided in Table 2. Control variants of the experiment were laid in a hormone-free MS environment.

**Table 2.**
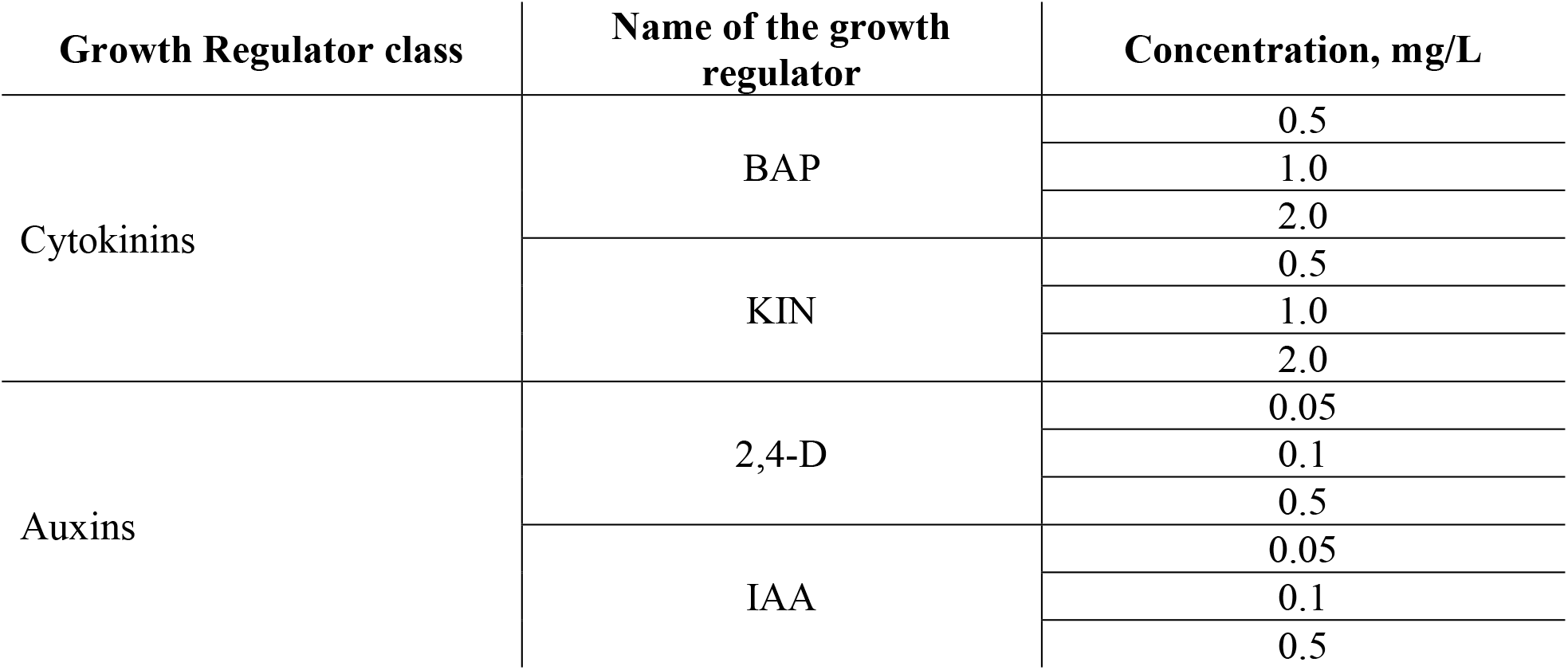
Variants of concentrations of growth regulators in modified MS media.

Micropropagation of shoots was carried out under aseptic conditions in a laminar flow box. Explants were planted in glass jars (5*7 cm), 3 in each. The distance between the shoots is 3-4 cm. The number of repetitions was sevenfold.

After inoculation of the shoots, their initial length was noted. After 7 and 14 days after planting, the following parameters were observed: change in shoot length (mm); rate of lateral shoot formation; shoot multiplication coefficient; frequency of rhizogenesis (%); number and length of roots (mm).

### Cultivation conditions, preparation of nutrient media

The seeds were sterilized in a laminar flow box using sterilized drying cabinet tools: scissors, tweezers. During the preparation of nutrient media, the pH was adjusted to 6.4 using a solution of potassium hydroxide with a concentration of 0.5 M. The sterilization of nutrient media was carried out in an autoclave MLS-3751 (Sanyo) at a temperature of 121 °C and a pressure of 1.5 atmospheres for 30 minutes. The seeds and shoots were cultivated at a constant temperature of 25 ° C and a sixteen-hour photoperiod.

### Statistical data processing

During the first part of the experiment, such parameters were determined as: seed germination – the ratio of the number of germinated seeds to the total number of seeds; the effectiveness of sterilization techniques, which was calculated by the ratio of the number of infected explants to the total number of explants planted.

In the second part of the experiment, the following parameters were monitored: the rate of regeneration of adventitious shoots is the percentage of the number of shoots with adventitious shoots from the total number of explants planted in culture (Guo et al., 2023); the frequency of rhizogenesis (%), which was determined by the proportion of shoots that successfully formed roots; the multiplication coefficient, as the average number of shoots formed on explants.

Statistical data processing was carried out using the Microsoft Excel program. A one-factor dispersion analysis of the following data was performed: the multiplication coefficient, an increase in the length of shoots, the length and number of roots formed; where the estimated influencing factors were selected: the type of nutrient medium, the type of growth regulator, and the concentration of growth regulators (Chokheli et al., 2023).

## RESULTS AND DISCUSSION

### Sterilization of explants

During the first part of the experiment, in vitro seeds were planted that were not subject to stratification, as a result of which it was revealed that methods No. I (C_2_H_5_OH (96%) + H_2_O_2_ (3%) in a 1:1 ratio with an exposure time of 7 minutes) and No.III (C_2_H_5_OH (70%) with an exposure time of 7 minutes) had the lowest contamination rates – 11%. The highest percentage of infected seeds was observed in method II (C_2_H_5_OH (96%) with an exposure time of 5 minutes) – 44% (Table 3).

**Table 3.** Estimate of the number of infected seeds that are not subject to cold stratification.

| Method number | I | II | III |
| --- | --- | --- | --- |
| % of infected seeds | 11% | 44% | 11% |

The result of the first part of the experiment showed that the most effective sterilization method was I (C_2_H_5_OH (96%) + H_2_O_2_ (3%) in a 1:1 ratio with an exposure time of 7 minutes) and III (C_2_H_5_OH (70%) with an exposure time of 7 minutes).

Observation for two weeks showed that the sprouts looked healthy, without any physiological abnormalities.

Similar results were obtained in Aicha (2013) studies when *T. satureioides* and *T.huemalis* were introduced into in vitro culture. Using a similar sterilization technique: 70% Ethanol with an exposure time of 3 minutes, followed by rinsing with 1% sodium hypochlorite solutions for 10 minutes and triple rinsing with distilled water, the percentage of contamination in the end did not exceed 5% in both species.

It was found that in the first part of the experiment, only those seeds that were sterilized by the first method overcame the dormancy stage. This may be due to the fact that a more aggressive sterilizing agent can facilitate the entry of nutrients from the nutrient medium into the embryo by chemically scarifying the seeds (Rosner et al., 2003). Because of this, only the First method of seed sterilization was used to sterilize seeds subject to emergency stratification.

In the second part of the experiment, the percentage of infected seeds was 2%. The sprouts also looked healthy, without any morphological abnormalities. The percentage of contamination in the second part of the experiment was significantly lower, which indicates the effectiveness of this technique.

### Germination of seeds in vitro

When seeds were planted *in vitro* without prior stratification, in two months only 8% of the seeds subjected to the first sterilization method were able to overcome the resting stage on the 1/2 MS medium.

In the second part of the experiment, germination of seeds subject to stratification occurred much faster (Figure 1): on the third day after planting on MS nutrient medium, seed germination was 32%, and on 1/2 MS and 1/4 MS 38.6% and 30.6%, respectively. After three more days, the germination rates were 41.3% (MS), 62.6% (1/2 MS), and 52% (1/4 MS). On day 9, the indicators remained almost unchanged and amounted to 44% (MS), 69.3% (1/2 MS) and 53.3% (1/4 MS).

**Figure 1.**
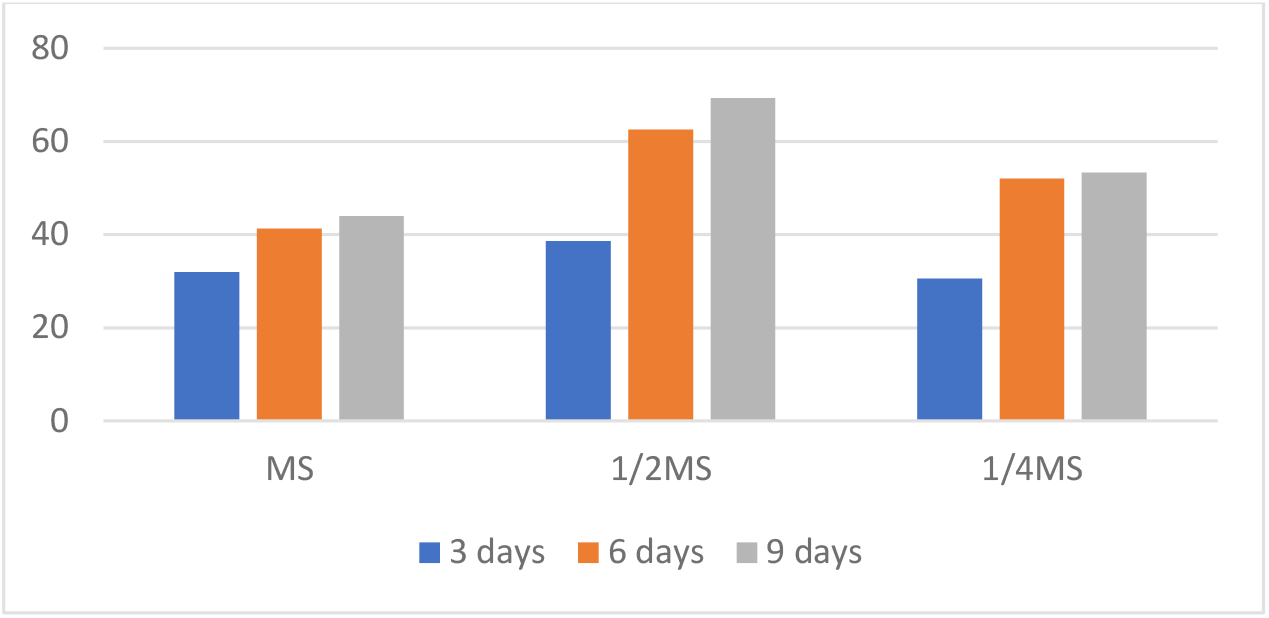
Germination rates of *T.calcareus* seeds in vitro on MS media for 9 days.

**Figure 2.**
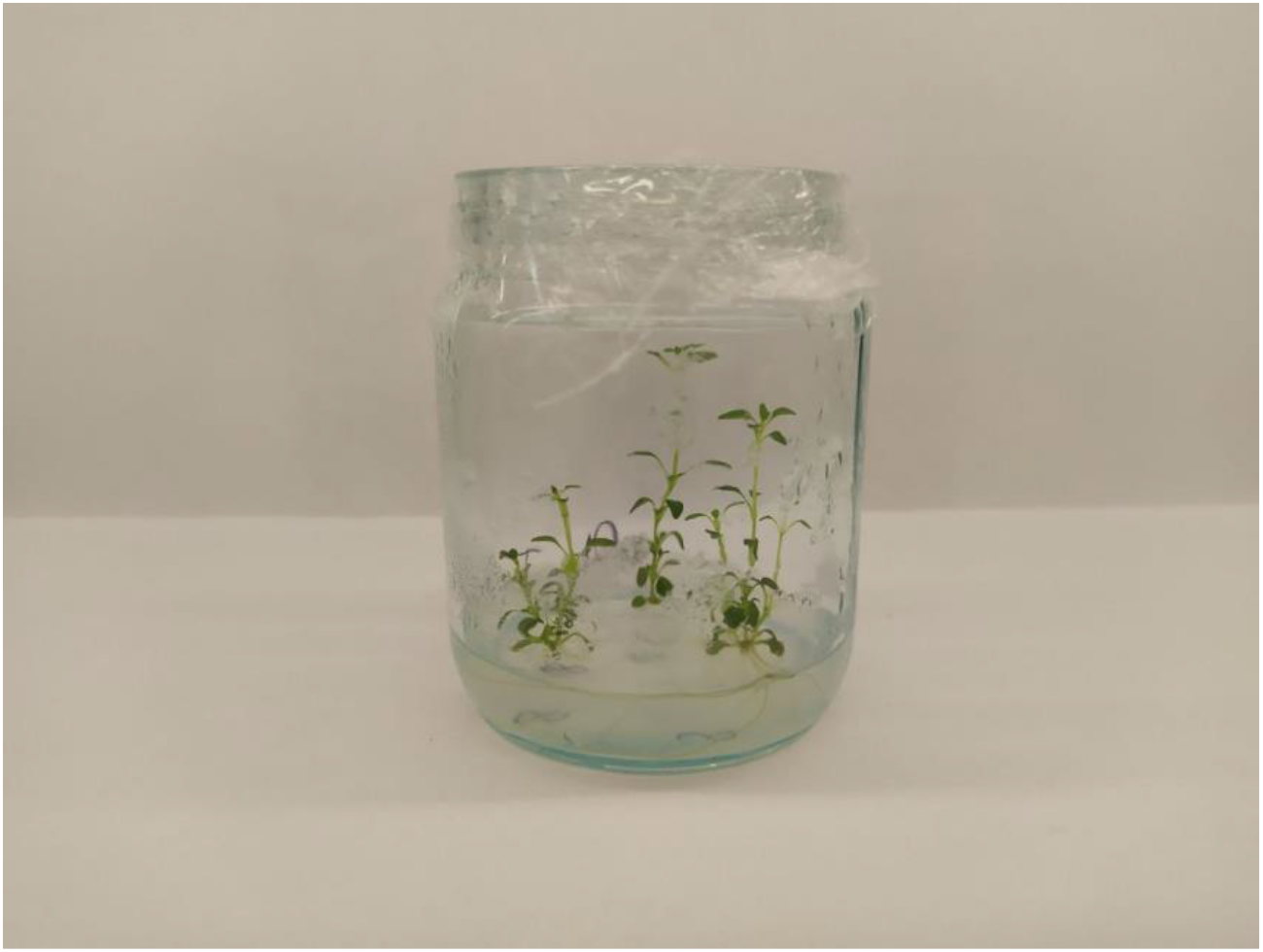
*T.calcareus* on the nutrient medium of 1/2 QL.

**Figure 3.**
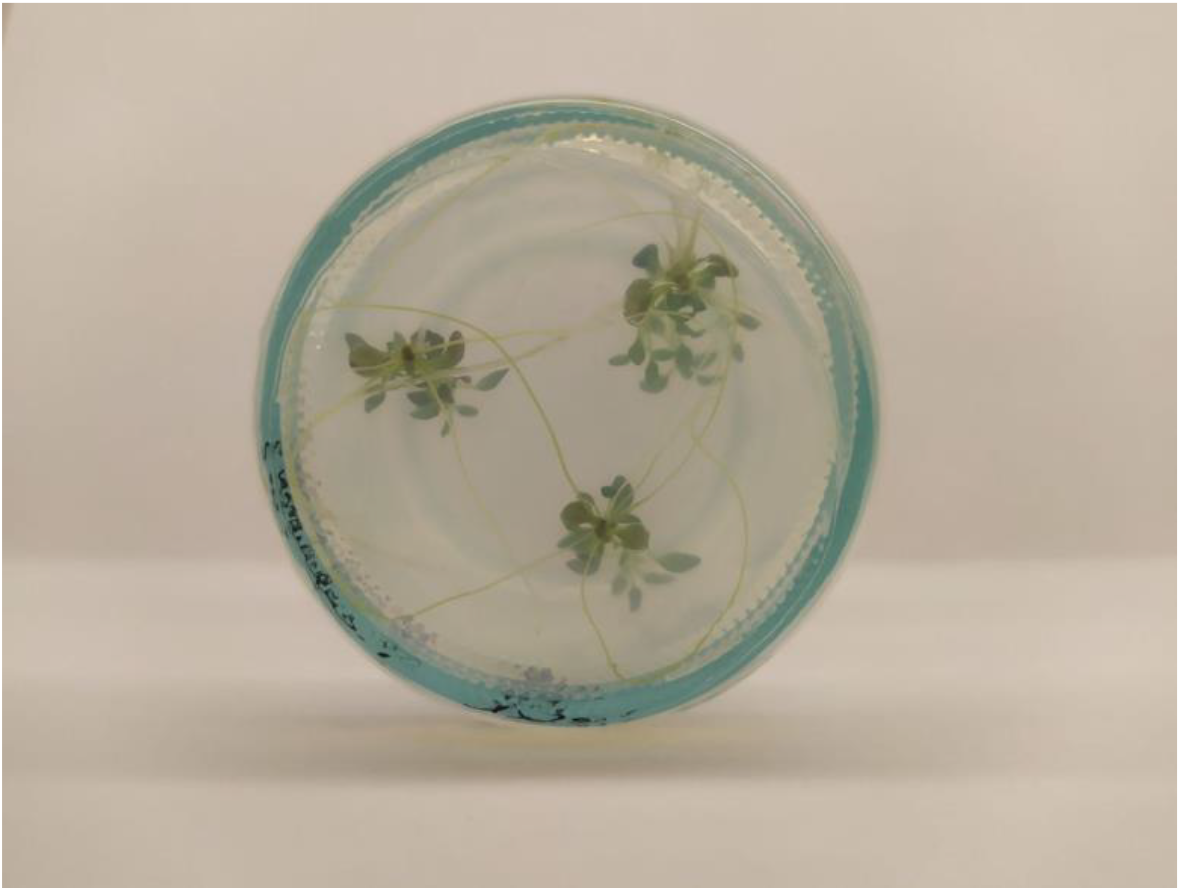
The roots of *T.calcareus* formed on the 1/2 QL media

During the experiment, it was also found that the sprouts on the 1/4 MS nutrient medium began to wither after a week and a half, while on the other media their growth took place without any deviations.

The seeds that were not stratified almost did not germinate (8%). In the second part of the experiment, there was a significant increase in the number of germinated seeds, as well as a decrease in their germination time, which indicates the need for stratification during germination of *T.calcareus* seeds.

As mentioned earlier, when germinating seeds in vitro of various species of the genus *Thymus*, many researchers do not subject the seed material to any stratification, for example, in *T. saturioides, T.huemalis* and *T.broussonetii*, the percentage of germinated seeds does not reach 35%, while in *T.vulgaris* it exceeded 60% (Affonco et al., 2009; Nordine et al., 2013a; Radomir et al., 2020). This may indicate that, depending on the place where the species grows, it is necessary to select its own stratification method for it.

The optimal medium for seed germination was 1/2 MS. This may be due to the fact that *T.calcareus* grows naturally in poor soils and excessive nutrient content can have an inhibitory effect on seed germination. Most studies of clonal reproduction of thyme when introduced into seed culture use only MS culture media as explants (Furmanova et al. 1992; Nordine et al., 2013b; Radomir et al., 2020). The observed difference between seed germination and their subsequent growth and development in this work suggests the need for a more careful selection of mineral substrates for seed germination.

### Multiplication and elongation of shoots in vitro

7 days after micropropagation, the following indicators were obtained: the multiplication coefficient, the average growth of shoots (mm) and the frequency of regeneration of lateral shoots (%). The results are presented in tables 4 and 5.

**Table 4.** Effect of various nutrients on multiplication coefficient, explant growth, and regeneration frequency of *T.calcareus in vitro*.

| Nutrient medium | The frequency of regeneration of lateral shoots, % | The growth of explants is average, mm | The multiplication coefficient is average |
| --- | --- | --- | --- |
| MS | 42.8 | 1.89±0.81 | 1.44±0.49 |
| 1/2 MS | 76.1 | 3.00±2.4 | 1.89±0.40 |
| B5 | 52.5 | 1.78±1.13 | 1.78±0.69 |
| 1/2 B5 | 33.3 | 1.22±0.96 | 1.44±0.59 |
| QL | 57.1 | 2.22±1.62 | 1.67±0.60 |
| 1/2 QL | 85.7 | 4.22±2.46 | 3.00±0.89 |

**Table 5.** Effect of various growth regulators on multiplication coefficient, explant growth, and regeneration frequency of *T.calcareus in vitro*.

|  | The frequency of regeneration of lateral shoots, % | Growth of explants, mm | The multiplication coefficient |
| --- | --- | --- | --- |
| MS | 42.8 | 1.89±0.81 | 1.44±0.49 |
| MS + BAP 0.5 mg/L | 76.1 | 0.91±0.76 | 2.5±1 |
| MS + BAP 1 mg/L | 66.6 | 0.83±0.55 | 2.17±0.89 |
| MS + BAP 2 mg/L | 23.8 | 0.33±0.5 | 1.25±0.38 |
| MS + KIN 0.5 mg/L | 100 | 1.75±1 | 3.75±0.67 |
| MS + KIN 1 mg/L | 85.7 | 0.66±0.56 | 2.83±0.89 |
| MS + KIN 2 mg/L | 85.7 | 0.83±0.42 | 2.33±0.67 |
| MS + IAA 0.05 mg/L | 57.1 | 2.16±1.56 | 2.5±1.3 |
| **MS + IAA 0.1 mg/L** | 66.6 | <u>2.25±1.25</u> | 2.58±1.4 |
| **MS + IAA 0.5 mg/L** | <u>100</u> | 1.83±1.3 | <u>3.67±0.72</u> |
| **MS + 2,4 D 0.05 mg/L** | 76.1 | <u>3.83±2.8</u> | 2.3±0.83 |
| **MS + 2,4 D 0.1 mg/L** | 71.4 | 1.67±1.44 | 2.16±0.72 |
| **MS + 2,4 D 0.5 mg/L** | 23.8 | 1.42±0.96 | 1.41±0.63 |

On the studied variants of nutrient media, the variation in the regeneration frequency of lateral shoots varies from 33.3% (1/2 B5) to 85.7 (1/2 QL); the growth of shoots from 1.22±0.96 (1/2 B5) to 4.22±2.46 mm (1/2 QL); the multiplication coefficient from 1.44±0.49 (MS) to 3.00±0.89 (1/2 QL).

As we can see, the highest values are observed on the 1/2 QL nutrient medium; 1/2 MS has the highest values after 1/2 QL; while 1/2 B5 has the lowest rates of lateral shoot regeneration and average explant growth.

It is also worth noting that with a decrease in the concentration of macro- and micronutrients in MS and QL nutrient media, large values of the studied parameters are observed, while in B5, on the contrary, the indicators decrease.

When using modified versions of the MS medium with the addition of different concentrations of growth regulators, the following variations in values are observed: the regeneration rate of lateral shoots varies from 23.8% (MS + 2,4 D 0.5 mg/L; MS + BAP 2 mg/L) to 100% (MS + KIN 0.5 mg/L; MS + IAA 0.5 mg/L); increase in explants from 0.66±0.56 (MS + KIN 1 mg/L) to 3.83±2.8 (MS + 2,4 D 0.05 mg/L); multiplication from 1.25±0.38 (MS + BAP 2 mg/L) to 3.75±0.67 (MS + KIN 0.5 mg/L).

An increase in the concentration of BAP in the MS nutrient medium leads to a decrease in all the studied parameters, yellowing of the leaves and darkening of the explant stems were observed. At a minimum concentration, compared with the control, the regeneration rates of lateral shoots increase by 33.3% and the average value of the multiplication coefficient.

With the addition of KIN, there is a decrease in such indicators as the regeneration frequency of lateral shoots and the regeneration coefficient, but despite this, these indicators are still among the highest among all the studied growth regulators. The increase in explants decreases with increasing concentration, approaching values similar to BAP. Compared to the control, the regeneration rate of lateral shoots increases by 57.2%, and the multiplication increases by more than two times, while the values of the explant increment are slightly lower. The appearance of the plants remained unchanged.

In contrast to the above-mentioned growth stimulators, an increase in the concentration of IAA leads to an increase in the regeneration rate of lateral shoots by 57.2% and the multiplication coefficient by more than two times. The minimum and intermediate concentrations had the third and second highest values of elongation of explants among all growth regulators. Comparison with the control showed that all values except for the growth of cuttings at maximum concentrations of IAA were higher.

Just as in the first two cases, an increase in the concentration of 2,4 D leads to a decrease in all indicators. Darkening and deformation of the apical leaves in the values of 0.1 and 0.5 mg/L was observed. In addition, a significant increase in the number of adventitious roots, abundantly pubescent with root hairs, was observed on some shoots. At the lowest concentration, the highest values of the studied parameters are observed, the increase in explants among all the studied samples was the largest (3.83±2.8 mm). Compared to the control, all values were higher.

The statistical calculation data of the effect of nutrient media, growth regulators and their concentration during the first week are provided in the table 6 and 7.

**Table 6.** Results of a one-sided dispersion analysis of the influence of types of mineral media, growth regulators and their concentration on the growth of implants.

| Effect | One-dimensional criterion of significance for the growth of explants, mm |  |  |  |  |
| --- | --- | --- | --- | --- | --- |
|  | SS | Grade | MS | F | p |
| Mineral base | 51.72 | 5 | 10.344 | 2.009 | 0.09 |
| Growth Regulator | 64.972 | 3 | 21.657 | 7.719* | 0.0000082 |
| Concentration of the growth regulator | 29.625 | 2 | 14.813 | 4.878* | 0.0089418 |

\* Values with confidence at $p=0.05$ .

**Table 7.**
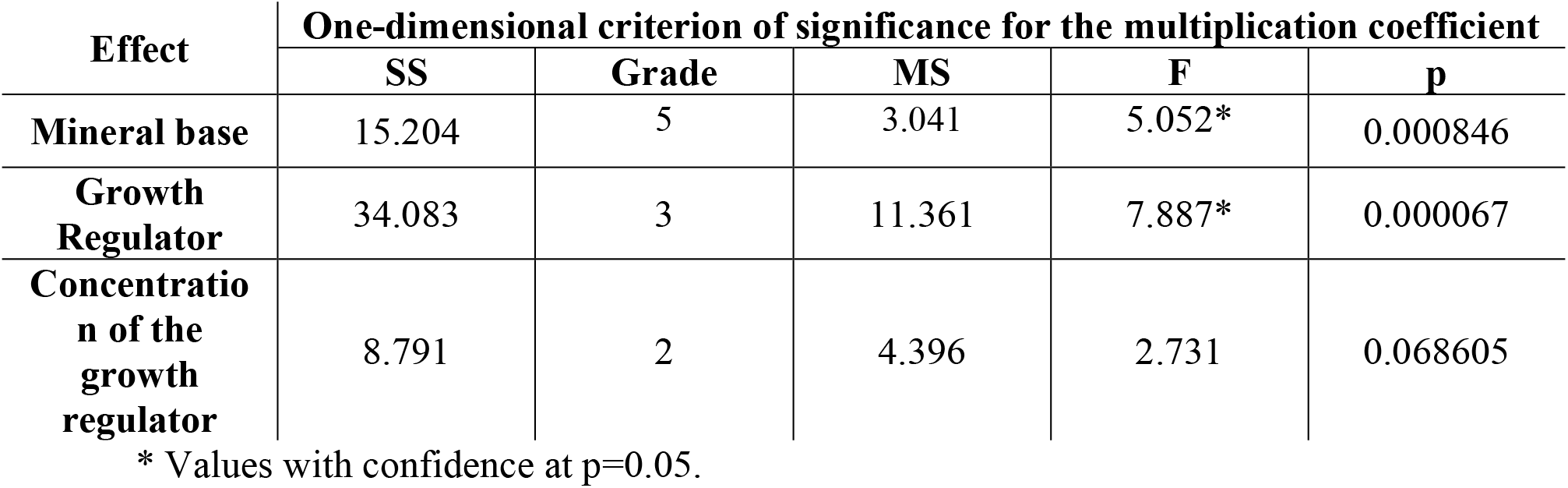
Results of a one-sided dispersion analysis of the influence of types of mineral media, growth regulators and their concentration on the multiplication coefficient.

| Effect | One-dimensional criterion of significance for the multiplication coefficient |  |  |  |  |
| --- | --- | --- | --- | --- | --- |
|  | SS | Grade | MS | F | p |
| Mineral base | 15.204 | 5 | 3.041 | 5.052* | 0.000846 |
| Growth Regulator | 34.083 | 3 | 11.361 | 7.887* | 0.000067 |
| Concentration of the growth regulator | 8.791 | 2 | 4.396 | 2.731 | 0.068605 |

\* Values with confidence at $p=0.05$ .

The mineral base does not significantly affect the elongation of explants. It was found that growth regulators and their concentration have a significant effect on the growth of explants. The F_fact_ values for the effectors were as follows: growth regulator – 7.71, growth regulator concentration – 4.87.

For the multiplication coefficient, a significant influence was found in the mineral base and growth regulator, with the following F_fac_ values: 5.05 and 7.88.

14 days after the start of the experiment, the following results were obtained for the coefficient of multiplication, elongation of shoots and the frequency of regeneration of lateral shoots (Table 8,9)

**Table 8.**
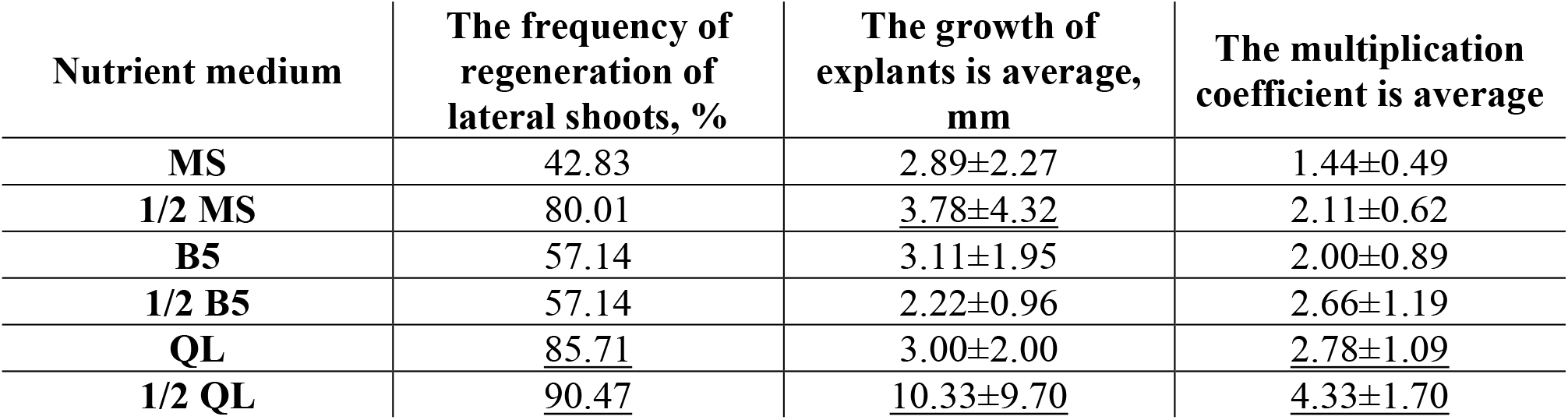
Effect of various nutrients on multiplication coefficient, explant growth, and regeneration frequency of *T.calcareus in vitro*.

**Table 9.** Effect of various regulators on multiplication coefficient, explant growth, and regeneration frequency of *T.calcareus in vitro*.

|  | The frequency of regeneration of lateral shoots, % | Growth of explants, mm | The multiplication coefficient |
| --- | --- | --- | --- |
| MS | 42.8 | <u>2.89±2.27</u> | 1.44±0.49 |
| MS + BAP 0.5 mg/L | 100 | 1.75±1.08 | 3.58±0.92 |
| MS + BAP 1 mg/L | 71.42 | 1.67±1.28 | 2.58±0.99 |
| MS + BAP 2 mg/L | 57.14 | 0.58±0.78 | 2.08±0.96 |
| MS + KIN 0.5 mg/L | <u>100</u> | 2.58±1.25 | <u>5.33±1.11</u> |
| MS + KIN 1 mg/L | <u>90.48</u> | 1.41±0.99 | <u>3.50±0.92</u> |
| MS + KIN 2 mg/L | <u>85.7</u> | 1.25±0.63 | 2.42±1.06 |
| MS + IAA 0.05 mg/L | 80.95 | <u>4.00±2.00</u> | 3.08±1.09 |
| MS + IAA 0.1 mg/L | <u>90.48</u> | 2.33±1.83 | 3.50±1.67 |
| MS + IAA 0.5 mg/L | <u>100</u> | 2.08±1.63 | <u>4.00±1.17</u> |
| MS + 2,4 D 0.05 mg/L | 76.19 | <u>5.08±2.31</u> | 2.83±1.06 |
| MS + 2,4 D 0.1 mg/L | 85.57 | 2.92±2.24 | 3.58±1.35 |
| MS + 2,4 D 0.5 mg/L | 47.61 | 1.75±1.38 | 1.50±0.58 |

There is an increase in the frequency of regeneration of lateral shoots (%) on all nutrient substrates except MS. The variation of the trait acquires large minimum and maximum values: 57.14% (B5, 1/2 B5) and 90.47% (1/2 QL). The increase in explants and the multiplication coefficient also have new variation values: 2.22±0.96 mm (1/2 B5) – 10.33±9.70 mm (1/2 QL); 1.44±0.49 (MS) – 4.33±1.70.

As before, an increase in all the studied values is observed on MS and QL media with a decrease in the concentration of micro- and macronutrients. Only on the B5 substrate is there a change in the trend of variability of the multiplication coefficient.

The variation in the regeneration frequency of lateral shoots after 14 days has a higher minimum value of 47.61% (MS + 2.4 D 0.5 mg/L). As with changes in the composition of nutrient media, the growth of explants and the multiplication coefficient now have new variation values: 0.58±0.78 mm (MS + BAP 2 mg/L) – 4.00±2.00 mm (MS + IAA 0.05 mg/L); 1.50±0.58 (MS + 2.4 D 0.5 mg/L) -5.33±1.11 (MS + KIN 0.5 mg/L).

The trend of changes in the studied parameters depending on the concentration of the growth regulator remains unchanged for BAP and KIN. The same is observed in IAA, with the exception of an increase in explants, with an increase in concentration, the elongation of shoots worsens. 2,4 D now has the highest regeneration rate of lateral shoots at a concentration of 0.1 mg/L.

There is yellowing of the leaves and darkening of part of the shoots at all concentrations of BAP, with an increase in the BAP content in the medium, the severity of these signs increases. In the case of KIN, the shoots of plants acquired a dark shade, but with a decrease in the level of growth stimulant in the medium, this effect weakened. A similar deviation was observed in some explants in environments with IAA.

On media with 2,4 D, a decrease and deformation of the apical leaves was periodically observed in all parameters of auxin concentration.

The data on the statistical calculation of the effect of nutrient media, growth regulators and their concentration after 14 days are provided in Tables 10 and 11.

**Table 10.** Results of a one-sided dispersion analysis of the influence of types of mineral media, growth regulators and their concentration on the growth of implants.

| Effect | One-dimensional criterion of significance for the growth of explants, mm |  |  |  |  |
| --- | --- | --- | --- | --- | --- |
|  | SS | Grade | MS | F | p |
| Mineral base | 414.44 | 5 | 82.899 | 2.742 | 0.0294037 |
| Growth Regulator | 100.18 | 3 | 50.090 | 10.396* | 0.0000613 |
| Concentration of the growth regulator | 82.354 | 2 | 27.4513 | 5.513* | 0.0013141 |

\* Values with confidence at $p=0.05$ .

**Table 11.** Results of a one-sided dispersion analysis of the influence of types of mineral media, growth regulators and their concentration on the multiplication coefficient.

| Effect | One-dimensional criterion of significance for the multiplication coefficient |  |  |  |  |
| --- | --- | --- | --- | --- | --- |
|  | SS | Grade | MS | F | p |
| **Mineral base** | 44,667 | 5 | 8,933 | 4,729 | 0,001362 |
| **Growth Regulator** | 36,167 | 3 | 18,083 | 7,825 | 0,000598 |
| **Concentration of the growth regulator** | 33,222 | 2 | 11,016 | 4,716 | 0,003629 |

\* Values with confidence at $p=0.05$ .

Two weeks later, there is an increase in the influence of all factors on the elongation of explants, so the value of F_fact_ increased for growth regulators and their concentrations by 2.68 and 0.64, respectively.

The null hypothesis for the mineral base has also been proven, which indicates an increase in its actual effect on elongation.

In the second week, the degree of influence on the multiplication coefficient slightly weakens for the mineral base and growth regulator, the F_fact_ values decrease by 0.33 and 0.06, respectively.

At the same time, the null theory for the concentration of the growth regulator is confirmed, the Ff_act_ of which is 4.71.

The multiplication coefficient, the growth of explants, and the regeneration frequency of *T.calcareus* lateral shoots had high variation values among all nutrient substrates. The 1/2 QL nutrient base proved to be the most effective for multiplication and elongation of explants with the following multiplication coefficient values – 4.33 ± 1.70, explant growth – 10.33 ± 9.70 mm and regeneration of lateral shoots – 90.47% (Table 8). This may be due to the fact that the QL nutrient substrate has the highest concentrations of calcium salts compared to MS and B5. The media also differ in the content of nitrogen salts, so B5 has the highest. Due to the fact that *T.calcareus* grows on Cretaceous limestone rock stands and on loose chalk with the onset of humus accumulation (Demina et al., 2016), media containing small amounts of nitrogen-containing compounds and organic compounds will be more favorable for its introduction into *in vitro*.

When studying the effect of growth regulators on the above-mentioned indicators, it was found that the regeneration rate of lateral shoots had the highest rates with the following modifications of the MS nutrient medium: MS + BAP 0.5 mg/L; MS + KIN 0.5 mg/L; MS + IAA 0.5 mg/L. The largest increase in explants (mm) was recorded at MS + 2.4 D 0.05 mg/L, and the multiplication at MS + KIN 0.5 mg/L.

When using BAP, a large number of explants showed chlorosis and darkening of stems, which indicates the unsuitability of this cytokinin for clonal reproduction of *T.calcareus*.

According to the calculations of the one-factor analysis of variance, it was revealed that the mineral base, the type of growth stimulant and its concentration have a significant effect on the elongation and multiplication of explants. Over time, the effect of all factors increases for the growth of shoots, to a greater extent for the growth regulator.

For the multiplication coefficient, the same effect is not observed in all factors except for the concentration of the growth regulator, but despite this, the F_fact_ is the largest for the growth regulator factor.

From this, it can be concluded that for multiplication and elongation of explants, it is most effective to use the 1/2 QL nutrient medium with the addition of small concentrations of KIN.

### Rhizogenesis of T.calcareus in vitro

During the experiment, when studying the effect of nutrient substrates and their modification on such indicators as: rhizogenesis %, root length mm and the average number of formed roots on the explant (pcs.). After 7 days, the following results were obtained (Tables 12 and 13).

**Table 12.** Effect of various nutrient media on the proportion of rhizogenesis, the average number of roots per explant, and the average root length of *T.calcareus in vitro*.

| **Nutrient medium** | **The frequency of rhizogenesis, %** | **Number of roots per explant, pcs.** | **The length of the roots, mm** |
| --- | --- | --- | --- |
| **MS** | 33.33 | <u>3.00±0.67</u> | 9.33±0.53 |
| **1/2 MS** | 52.38 | 2.00±1.00 | 6.38±0.60 |
| **B5** | 42.85 | 1.75±0.75 | 9.43±0.83 |
| **1/2 B5** | 42.85 | 1.5±0.5 | 7.33±0.73 |
| **QL** | <u>66.66</u> | 2.67±0.89 | <u>10.31±0.71</u> |
| **1/2 QL** | <u>90.47</u> | <u>3.5±1.5</u> | <u>12.96±1.12</u> |

**Table 13.** The effect of various growth regulators and their concentrations on the proportion of rhizogenesis, the average number of roots per explant, and the average root length of *T.calcareus in vitro*.

|  | The frequency of rhizogenesis, % | Number of roots per explant, pcs. | The length of the roots, mm |
| --- | --- | --- | --- |
| MS | 33.33 | $3.00 \pm 0.67$ | $9.33 \pm 0.53$ |
| MS + BAP 0.5 mg/L | 19.04 | $1.5 \pm 0.5$ | $5.00 \pm 3.61$ |
| MS + BAP 1 mg/L | – | – | – |
| MS + BAP 2 mg/L | 9.52 | $1.00 \pm 0.00$ | $3.00 \pm 0.00$ |
| MS + KIN 0.5 mg/L | 23.80 | $1.67 \pm 0.44$ | $10.20 \pm 2.86$ |
| MS + KIN 1 mg/L | – | – | – |
| MS + KIN 2 mg/L | 14.28 | $1.00 \pm 0.00$ | $5.5 \pm 4.96$ |
| MS + IAA 0.05 mg/L | <u>66.66</u> | $2.00 \pm 0.75$ | $4.25 \pm 1.73$ |
| MS + IAA 0.1 mg/L | 33.33 | $2.25 \pm 0.88$ | $3.00 \pm 1.73$ |
| MS + IAA 0.5 mg/L | 28.57 | <u>$4.50 \pm 2.25$</u> | $5.67 \pm 3.14$ |
| MS + 2,4 D 0.05 mg/L | <u>57.12</u> | $2.57 \pm 0.78$ | $2.89 \pm 1.08$ |
| MS + 2,4 D 0.1 mg/L | – | – | – |
| MS + 2,4 D 0.5 mg/L | – | – | – |

On the studied variants of nutrient media, the frequency of rhizogenesis varies from 33.33% (MS) to 90.47% (1/2 QL), the average number of roots per explant from 1.5±0.5 (1/2 B5) to 3.5±1.5 (1/2 QL), and the average root length from 6.38±0.6 (1/2 MS) to 12.96±1.12 (1/2 QL).

As with the multiplication of shoots, the highest rates are observed on a nutrient medium of 1/2 QL.

With a decrease in the concentration of macro- and micronutrients, the frequency of rhizogenesis increases in all media except B5, where it remains unchanged; the average number of roots per explant, in turn, decreases in all media except QL, the same trend is observed with the average root length.

When using modified versions of the MS medium with the addition of various concentrations of growth stimulants, the following variations are observed: the proportion of rhizogenesis varies from 9.52% (MS+BAP 2 mg/L) to 66.66% (MS +IAA 0.05 mg/L); the average number of roots per explant is from 1.00± 0.0 (MS+BAP 2 mg/L; MS+KIN 2 mg/L) to 4.50±2.25 (MS+IAA 0.05 mg/L); average root length from 2.89±1.08 (MS+2.4 D 0.05 mg/L) to 10.20±2.86. (MS+KIN 0.5 mg/L).

With a decrease in cytokinin concentration, an increase in all indicators is observed. When compared with the control, all values for BAP are lower, and almost the same is observed for KIN, except for the length of the roots.

With an increase in the concentration of IAA, the proportion of rhizogenesis decreases, the number of roots on the explant increases, but there is no effect on the length of the roots.

An increase in the concentration of 2,4 D significantly inhibits rhizogenesis, and intensive callus formation was observed on some explants. It was also found that at minimal concentrations of this auxin, adventitious roots, abundantly pubescent with root hairs, began to form.

Statistical calculation data on the effect of nutrient media, growth regulators and their concentration on rhizogenesis 7 days after explant inoculation are provided in Tables 14 and 15.

**Table 14.** Results of a one-sided dispersion analysis of the effect of types of mineral media, growth regulators and their concentration on root length.

| Effect | One-dimensional significance criterion for the length of the roots on the explant. |  |  |  |  |
| --- | --- | --- | --- | --- | --- |
|  | SS | Grade | MS | F | p |
| Mineral base | 283.736 | 5 | 56.747 | 1.489 | 0.206114 |
| Growth Regulator | 179.553 | 3 | 59.851 | 9.688* | 0.000020 |
| Concentration of the growth regulator | 42.128 | 2 | 21.064 | 2.607 | 0.081010 |

\* Values with confidence at $p=0.05$ .

**Table 15.**
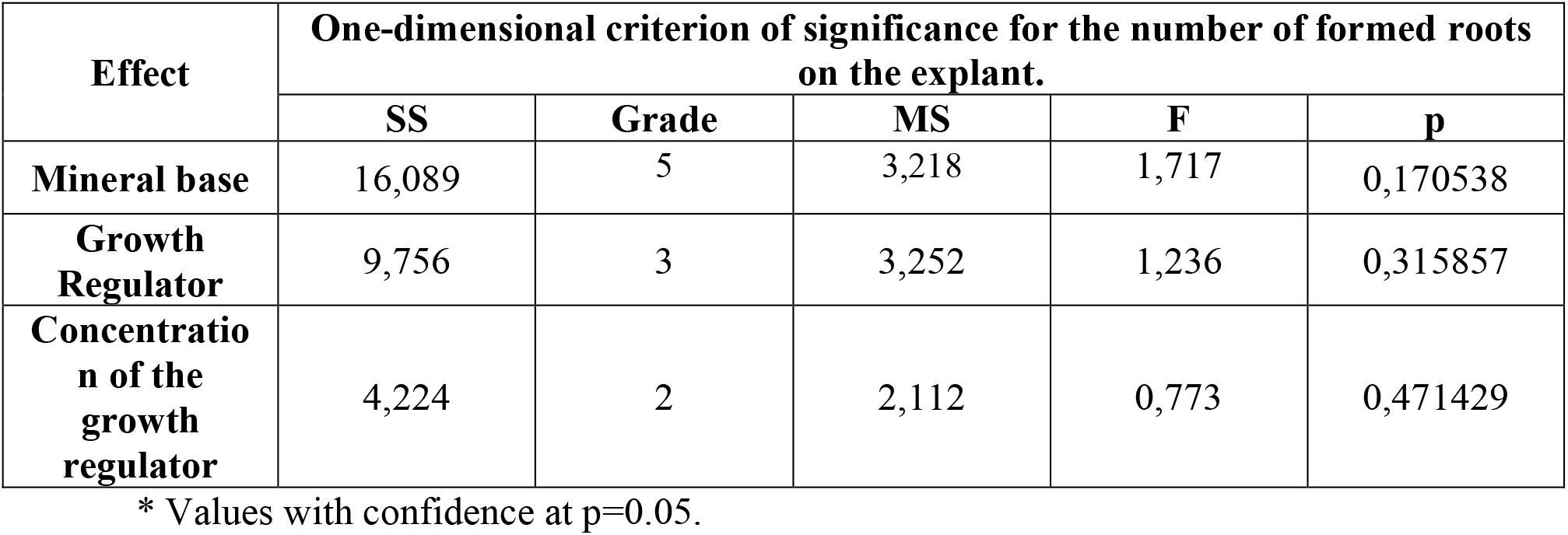
Results of a one-sided dispersion analysis of the influence of types of mineral media, growth regulators and their concentration on the number of formed roots on the explant.

| Effect | One-dimensional criterion of significance for the number of formed roots on the explant. |  |  |  |  |
| --- | --- | --- | --- | --- | --- |
|  | SS | Grade | MS | F | p |
| Mineral base | 16,089 | 5 | 3,218 | 1,717 | 0,170538 |
| Growth Regulator | 9,756 | 3 | 3,252 | 1,236 | 0,315857 |
| Concentration of the growth regulator | 4,224 | 2 | 2,112 | 0,773 | 0,471429 |

\* Values with confidence at $p=0.05$ .

For root length, a significant effect was found for the growth regulator, where F_fact_ was 9.688.

After 14 days, the following indicators of the frequency of rhizogenesis were obtained, cf. the number of roots per explant and cf. the length of the roots (Tables 16 and 17).

**Table 16.** Effect of various nutrient media on the proportion of rhizogenesis, the average number of roots per explant, and the average root length of *T.calcareus in vitro*.

| Nutrient medium | The frequency of rhizogenesis, % | Number of roots per explant, pcs. | The length of the roots, mm |
| --- | --- | --- | --- |
| MS | 52.38 | 2.40±0.88 | 16.58±6.18 |
| 1/2 MS | 57.14 | 2.60±0.72 | 15.23±6.28 |
| B5 | 57.14 | 2.00±0.08 | 17.80±6.80 |
| 1/2 B5 | 57.14 | 2.60±0.72 | 13.85±8.58 |
| QL | 66.66 | 2.83±1.11 | 23.41±8.46 |
| 1/2 QL | 100 | 3.00±1.33 | 28.41±11.96 |

**Table 17.** Effect of various growth regulators and their concentrations on multiplication coefficient, explant growth, and regeneration frequency of *T.calcareus in vitro*.

|  | The frequency of rhizogenesis, % | Number of roots per explant, pcs. | The length of the roots, mm |
| --- | --- | --- | --- |
| MS | 52.38 | 2.40±0.88 | 16.58±6.18 |
| MS + BAP 0.5 mg/L | 19.04 | 1.5±0.50 | 5.00±3.61 |
| MS + BAP 1 mg/L | — | — | — |
| MS + BAP 2 mg/L | 9.52 | 1.00±0.00 | 7.00±0.00 |
| MS + KIN 0.5 mg/L | 23.80 | 1.75±0.39 | 12.14±2.16 |
| MS + KIN 1 mg/L | — | — | — |
| MS + KIN 2 mg/L | 14.28 | 1.00±0.00 | 7.50±3.50 |
| MS + IAA 0.05 mg/L | 66.67 | 3.00±1.14 | 11.05±6.53 |
| MS + IAA 0.1 mg/L | 42.85 | 3.00±0.80 | 6.53±2.36 |
| MS + IAA 0.5 mg/L | 38.10 | 5.25±1.88 | 11.23±3.41 |
| MS + 2,4 D 0.05 mg/L | 58.33 | 3.60±0.64 | 3.33±1.41 |
| MS + 2,4 D 0.1 mg/L | — | — | — |
| MS + 2,4 D 0.5 mg/L | — | — | — |

On the 14th day of observations, the proportion of rhizogenesis ranged from 52.38% (MS) to 100% (1/2 QL). The variation in the average number of roots per explant was reduced from 2.00±0.08 (B5) to 3.00±1.33 (1/2 QL), due to the fact that explants in which rhizogenesis had only recently been initiated had a small number of them. The variation in root length ranged from 13.85±8.58 (1/2 B5) to 28.41±11.96 (1/2 QL).

The highest rates of rhizogenesis, number of roots per explant, and root length are now observed only in QL media.

As in the early observations, a decrease in the concentration of macro- and micronutrients leads to an increase in the proportion of rhizogenesis in all media except B5. Also, large root lengths are now observed on full MS and B5 environments, while the opposite is true for QL.

The variation in the values of the rhizogenesis fraction on the modifications of the MS medium after 14 days was unchanged. The maximum variation in the average number of roots per explant was also observed on MS+IAA 0.5 mg/L medium and was 5.25±1.88 pcs. The minimum root length values were also observed on the same medium throughout the experiment: MS+2.4 D 0.05 mg/L with values of 3.33±1.41 mm.

As before, an increase in all indicators is recorded with a decrease in cytokinin concentration values. All the studied indicators on the BAP and KIN media are less than controlled.

The effect of IAA and 2,4 D on rhizogenesis, length, and number of roots remained unchanged. When compared with the control, both auxins at minimal concentrations had higher values of the proportion of rhizogenesis.

Statistical calculation data on the effect of nutrient media, growth regulators and their concentration on rhizogenesis 14 days after explant inoculation are provided in Tables 18 and 19.

**Table 18.** Results of a one-sided dispersion analysis of the effect of types of mineral media, growth regulators and their concentration on root length.

| Effect | One-dimensional criterion of significance for the number of formed roots on the explant. |  |  |  |  |
| --- | --- | --- | --- | --- | --- |
|  | SS | Grade | MS | F | p |
| Mineral base | 3.738 | 5 | 0.748 | 0.447 | 0.811430 |
| Growth Regulator | 38.456 | 2 | 12.819 | 5.883 | 0.003021 |
| Concentration of the growth regulator | 3.146 | 3 | 1.572 | 0.473 | 0.627813 |

\* Values with confidence at $p=0.05$ .

**Table 19.** Results of a one-sided dispersion analysis of the effect of types of mineral media, growth regulators and their concentration on the number of formed roots on the explant.

| Effect | One-dimensional significance criterion for the length of the roots on the explant. |  |  |  |  |
| --- | --- | --- | --- | --- | --- |
|  | SS | Grade | MS | F | p |
| Mineral base | 3013,282 | 5 | 602,656 | 5,077* | 0,000398 |
| Growth Regulator | 648,377 | 3 | 216,126 | 11,697* | 0,000000 |
| Concentration of the growth regulator | 51,152 | 2 | 25,576 | 1,138 | 0,325502 |

\* Values with confidence at $p=0.05$ .

The significant effect of the growth regulator on root length decreased after 14 days. The value for the week F_fact_ by 3.8 and amounted to 5.88.

There is a sharp increase in the importance of the mineral environment factor and growth regulator on the number of roots on the explant at p=0.05. The F_fact_ for these effectors was equal to: 5.07 and 11.69. The effect of the growth regulator is significantly higher than for the mineral base.

As with the indicators of multiplication and elongation of shoots, the parameters of rhizogenesis on different nutrient media differed significantly. The highest values were observed on the 1/2 QL nutrient medium, where the rhizogenesis fraction reached 100%, the average root length was 28.41±11.96 mm, and the number of roots per explant was 3.00±1.33. A decrease in the concentration of micro- and macronutrients had a positive effect on this indicator in all media except B5, where the values did not differ.

As mentioned earlier, the distinctive characteristics on QL media may be due to their closer chemical compositions to the soils of natural habitats.

When using modified MS growth regulators, the proportion of rhizogenesis was not so high and reached a maximum at MS+IAA 0.05 mg/L, where all the studied characteristics except root length exceeded the control. The root lengthening indices did not stand out so much, unlike in hormone-free nutrient media. The only parameter that significantly exceeded the control was the number of roots per explant. The maximum value of this indicator was observed at MS+IAA 0.5 mg/L.

An increase in the concentration of all growth regulators inhibits rhizogenesis, especially well manifested in 2,4 D, where callus neoplasms prevented rooting.

According to statistical calculations, it was revealed that only the growth regulator has a significant effect on the length of the roots, which weakens over time. The number of roots on the explant is significantly influenced by the composition of the nutrient medium and the growth regulator, where the F_fact_ of the growth regulator is significantly higher than that of the substrate.

In total, for the development of the root system of *T.calcareus in vitro*, it is recommended to use an 1/2 QL nutrient medium with low concentrations of IAA.

## CONCLUSION

Thus, in the course of this study, a protocol was developed for depositing a representative of the genus *Thymus*: *T. calcareus*. Protocols have been developed for the sterilization of primary explants (seeds); initiation (germination from seeds); multiplication and rhizogenesis *in vitro*. The next step in experiments on growing *T. calcareus* will be the development of callousogenesis protocols, followed by the transfer of calluses to suspension culture. This will help to obtain various secondary metabolites that make up thyme, which in the future will allow you to work in the pharmaceutical field. The last stage of cultivation is *T. calcareus*, in the future, will be the development of protocols for post-aseptic adaptation of plants, with the aim of further acclimatization in open ground conditions.

